# Low Level Whole-Brain Radiation Enhances Theranostic Potential Of Single Domain Antibody Fragments For HER2-Positive Brain Metastases

**DOI:** 10.1101/2022.04.19.488555

**Authors:** Daniele Procissi, Stephen A. Jannetti, Markella Zannikou, Zhengyuan Zhou, Darryl McDougald, Deepak Kanojia, Ganesan Vaidyanathan, Michael R. Zalutsky, Irina V. Balyasnikova

**Author notes:** These authors contributed equally (Daniele Procissi, Stephen A. Jannetti). Evergreen Theragnostics, Springfield, NJ 07081 USA. **Corresponding Author:** Irina V. Balyasnikova, Ph.D., Department of Neurological Surgery, Northwestern University, Chicago, IL 60611, USA. **Co-Corresponding Author:** Michael R. Zalutsky, Ph.D., Department of Radiology, Duke University Medical Center, 311 Research Drive, Durham, NC 27710, USA. Authorship statement: Study design: M.R.Z. and I.V.B. MRI data collection and analysis: D.P. Radiochemistry: Z.Z., D.M., and G.V. PET imaging data collection and analysis: Z.Z and S.A.J. Histology, Flow Cytometry and Western Blot: I.V.B., M.Z, D.K. Final manuscript review: I.V.B. and M.R.Z.

## Abstract

**Background:** Single-domain antibody fragments (aka V_H_H, ∼13 kDa) are promising delivery systems for brain tumor theranostics; however, achieving efficient delivery of V_H_H to intracranial lesions remains challenging due to the tumor-brain barrier. Here, we evaluate low-dose whole-brain irradiation as a strategy to increase the delivery of an anti-HER2 V_H_H to breast cancer-derived intracranial tumors in mice.

**Methods:** Mice with intracranial HER2-positive BT474BrM3 tumors received 10-Gy fractionated cranial irradiation and evaluated using non-invasive imaging methods. The anti-HER2 V_H_H 5F7 was labeled with ^18^F, administered intravenously to irradiated mice and controls, and PET/CT imaging was conducted at various intervals after irradiation. Tumor uptake of _18_F-labeled 5F7 in irradiated and control mice was compared by PET/CT image analysis and correlated with tumor volumes. In addition, longitudinal dynamic contrast-enhanced MRI (DCE-MRI) was conducted to visualize and quantify the potential effects of radiation on tumor perfusion and permeability.

**Results:** Increased ^18^F-labeled 5F7 intracranial tumor uptake was observed with PET in mice that received cranial irradiation, with maximum tumor accumulation seen approximately 12 days post initial radiation treatment. No radiation-induced changes in HER2 expression were detected by Western blot, flow cytometry, or on tissue sections. DCE-MRI imaging demonstrated transiently increased tumor perfusion and permeability after irradiation, consistent with the higher tumor uptake of _18_F-labeled anti-HER2 5F7 in irradiated mice.

**Conclusion:** Low-level brain irradiation induces dynamic changes in tumor vasculature that increase the intracranial tumor delivery of an anti-HER2 V_H_H, which could facilitate the use of labeled sdAb to detect, monitor, and treat HER2-expressing brain metastases.

**Key points:**

- Low-level radiation enhances uptake of HER2-specific V_H_H in intracranial tumors.
- XRT + radiolabeled V_H_H shows promise as a treatment strategy for breast cancer brain metastases.

**Importance of the Study:** Improving the detection and treatment of brain metastases (BM) that overexpress human epidermal growth factor receptor type 2 (HER2) is an urgent medical need. Drug delivery to BM is confounded by their tumor vasculature, which is more restrictive than in GBM. Single domain antibody fragments, about one-tenth the size of antibodies, could be promising theranostic vectors for BM provided sufficient BM uptake could be achieved. In this study, we utilized longitudinal PET imaging to demonstrate that low-dose whole-brain irradiation (WBRT) significantly increased ^18^F-labeled HER2-specific 5F7 V_H_H uptake in intracranial HER2-positive tumors in mice. Combining low dose WBRT with 5F7 V_H_H labeled with α-or β-particle emitting radionuclides could provide an effective and specific targeted radiotherapeutic strategy for treating patients with HER2-expressing BM.

## Introduction

Breast cancer affects approximately one in eight women and is the second leading cause of cancer death in women in the United States.^1^ Breast cancers are heterogeneous, with subtypes often dictating where metastases are most likely to arise. These subtypes are associated with different survival characteristics and help to inform surveillance and treatment options.^2^ While systemic treatment has increased survival for metastatic breast cancer patients, the development of central nervous system (CNS) metastases in breast cancer patients is on the rise leading to a poor prognosis.^3-6^ The overall survival of patients with brain metastases of breast cancer (BCBM) is crucially dependent on the early diagnosis of CNS involvement.^7^ HER2 overexpression is found in approximately 30% of all breast cancer cases.^8,9^ In HER2-positive patients, the rate of CNS metastases ranges from 30 to 50%,^10-12^ making HER2 an attractive target for the detection and treatment of BCBM.

In patients with BCBM, the blood-tumor barrier (BTB) hinders the delivery of clinically effective treatments. Despite being physically and heterogeneously impaired to a varying degree in almost all brain metastases, the BTB still effectively prevents significant tumor penetration of most chemotherapeutics and imaging agents in over 90% of cases.^13^ The inability to reach the tumor tissue thus limits both the therapeutic and diagnostic efficacy of drug/agent-based approaches.

Whole-brain radiation therapy (WBRT) remains one of the most widely utilized treatment strategies for BCBM because it circumvents the BTB by providing a therapeutic effect without targeting tumor tissue via an exogenously administered agent.^14,15^ WBRT, in conjunction with stereotactic radiosurgery, can provide a survival benefit, albeit only when treating single brain metastasis. The WBRT protocol consisted of 2.5 Gy daily over two weeks (37.5 Gy total), a dose known to increase the permeability of the BBB in human patients diagnosed with brain metastases.^16^

The HER2-targeted monoclonal antibody (mAb) trastuzumab has permitted PET imaging of BCBM in patients;^17^ however, generally only in those with significant BBB disruption resultant from a relatively high dose of WBRT or advanced tumors. A promising strategy for circumventing this limitation is the development of smaller HER2-targeted scaffolds for PET imaging and radiopharmaceutical therapy, such as single-domain antibody fragments (aka V_H_H or nanobodies). V_H_H are derived from Camelids, have a molecular weight of 12-15 kDa, have low immunogenicity, and can have a subnanomolar affinity.^18,19^ Importantly, studies in murine models with anti-HER2 V_H_H have demonstrated considerably more rapid tumor penetration of V_H_H compared with mAbs and the ability to localize in HER2-positive intracranial xenografts.^20^ Moreover, the feasibility of imaging a patient with brain metastases using an anti-HER2 V_H_H has been reported.^21^

The current study utilizes the anti-HER2 V_H_H 5F7^22^ to investigate the effects of low-level WBRT on its tumor uptake in an orthotopic BCBM mouse model using a non-invasive imaging approaches. We hypothesize that by using WBRT fractions below 3 Gy, V_H_H-based imaging agents and therapeutics could be made more effective without the quality of life impairments usually associated with higher radiation doses.^23^ Importantly, the proposed strategy would enable clinicians to track transient changes in tumor physiology, including permeability, to identify optimal therapeutic and diagnostic windows.^16^

## Methods

### General

The primary antibodies utilized for protein analysis by western blot were rabbit anti-human HER2/ERBB2 (D8F12), rabbit anti-human GAPDH (Cell Signaling Technologies, MA), and secondary goat anti-rabbit IgG conjugated with horseradish peroxidase (HRP) (Thermo Fisher Scientific, IL). To detect cell-surface HER2 expression by flow cytometry, we used Herceptin^®^ at 5 µg/ml and goat anti-human Alexa Fluor^®^ 647 (Sigma, St. Louis, MO) at 1:750 dilution. Human IgG1 served as a negative control. For HER2 staining of tissue sections, we utilized rabbit monoclonal antibody clone 29D) (Cell Signaling Technologies) at 1:200 dilution, followed by detection with anti-rabbit IgG conjugated to HRP.

### Cell Culture

HER2-overexpressing BT474BrM3 cells^24^ (hereafter referred to as BT474Br) were originally obtained from Dr. D. Yu (The University of Texas, MD Anderson Cancer Center) and modified via lentiviral transduction to express either firefly luciferase (fluc) or both fluc and mCherry fluorescent protein. Cells were maintained in DMEM media (Corning) supplemented with 10% FBS (HyClone, UT) and penicillin/streptomycin. BT474Br cells were tested for mouse pathogens before using them in animal experiments. In addition, modified BT474Br cells also were routinely tested for mycoplasma.

### Animal Studies

All animal studies were performed under protocols approved by the Institutional Animal Care and Use Committees at Northwestern University and Duke University. Six to eight-week-old female athymic nude mice were obtained from Jackson laboratories. Intracranial xenografts were established by injecting 4×10^5^ BT474Brfluc cells in the right frontal lobe using a Hamilton syringe with a 26-gauge needle.^25^ Following implantation, tumor growth was monitored by bioluminescent imaging.

### Radiation Therapy

Mice with BT474Br intracranial xenografts were divided into control and irradiated groups. Mice in the irradiated group underwent WBRT for five consecutive daily 2-Gy doses using a GammaCell irradiator as previously described.^26^ Specifically, mice were anesthetized with ketamine/xylazine, and their bodies were shielded with lead except for the cranium.

### Radiolabeling 5F7 Anti-HER2 V_H_H

5F7^22^ was labeled with _18_F using the *N*-succinimidyl ester-containing residualizing prosthetic agent RL-III.^27^ Briefly, a solution of 5F7 in borate buffer, pH 8.5 was incubated with [_18_F]RL-III at room temperature for 20 min, and the resultant [_18_F]RL-III-5F7 conjugate was isolated by gel filtration over a PD10 column (GE Healthcare, Piscataway, NJ) eluted with PBS. Radiochemical purity, HER2-binding affinity, and immunoreactivity of [_18_F]RL-III-5F7 (hereafter referred to as _18_F-5F7) were determined as reported previously.^27^

### PET Imaging

PET/CT imaging was performed on a Siemens Inveon micro-PET/CT system (Malvern, PA). Mice were imaged 30 min post-i.v. injection of 1.5 ± 0.4 MBq (18.4 ± 10.9 µg) of _18_F-5F7. Mice were anesthetized using 2–3% isoflurane in oxygen and placed prone in the scanner gantry for a 5 min static PET acquisition followed by a 5 min CT scan. List mode PET data were histogram-processed, and images were reconstructed using a standard OSEM3D/MAP algorithm — 2 OSEM3D iterations and 18 MAP iterations — with a cutoff (Nyquist) of 0.5. Images were corrected for attenuation (CT-based) and radioactive decay. Images were analyzed using Inveon Research Workplace software.

### Histology

Following PET imaging, mice were euthanized, and brains were removed and placed in buffered 10% formalin. Paraffin blocks were prepared for each animal and sectioned as 4µm tissue sections. Tissue sections were processed for either hematoxylin and eosin staining (H&E) or HER2 expression. For this, tissue sections were deparaffinized, and antigen retrieval was performed using citrate buffer pH 6 at 110°C for 20 min in a pressure cooker. Sections were blocked with 5% goat serum in tris-buffered saline and 0.1% Tween® 20 detergent (TBST) for 30 min and stained with anti-HER2 antibodies at 1:500 dilution (Cell Signaling #21650). After three washes with TBST, the primary antibody was detected with Mach2 rabbit HRP-Polymer (Biocare Medical, Concord, CA) followed by incubation with DAB substrate for 5 min. HER2 expression was evaluated in four tissue sections from three brains in both the control and irradiated groups. To determine tumor volume, 30 slides (with four tissue sections placed on each slide) were prepared from each brain, and every fifth slide was stained for H&E. Stained sections were scanned and analyzed in NDP.view 2.8.24 (Hamamatsu Photonics K.K.). The tumor area was identified and traced with the cursor to obtain area measurements. Volumetric analysis was conducted using the total tumor area per section multiplied by the thickness of sections and the number of slices collected from each brain.

### Flow Cytometry

BT474Br cells harvested from the mouse intracranial tumors were stained with anti-HER2 antibody (Herceptin_®_) followed by detection with anti-human IgG conjugated with Alexa Fluor^®^ 647. Isotype-matched human IgG1 served as a negative control. Flow cytometry analysis was performed using a BD FACSCalibur instrument (BD Biosciences; San Jose, CA). Data were collected as both percent positive cells and median fluorescence intensity (MFI).

### Western Blot

For analysis of HER2 expression, BT474Br cells were harvested from control and irradiated mice and plated for 2-3 days in culture to remove contaminating cells. Adherent BT474Br cells were then harvested and lysed for protein analysis using a standard western blot technique. Membranes with their transferred proteins were blocked with TBS-Tween20 buffer supplemented with 1% BSA and then stained with anti-HER2 or GAPDH antibodies at 1:2000 and 1:5000 dilutions, respectively. After 3 washes with TBS-Tween20 buffer, primary antibodies were detected with 1:5000 anti-rabbit antibody-HRP conjugate. Unbound antibodies were washed and identified using the Calrity™ Western ECL substrate (Bio-Rad, Hercules, CA). Densitometry was done using ImageJ software (NIH). HER2 expression was calculated as the ratio of HER2 to GAPDH in each sample.

### MRI and DCE-MRI Imaging

MRI was conducted on a 7T Bruker ClinScan. After placing a tail vein catheter for delivery of gadolinium (Gd_3+_) contrast (average dose ∼0.01 mmol/kg), each mouse was anesthetized using a mixture of O_2_/100% and isoflurane and then placed in the scanner. Brain MR images were acquired using a dedicated brain 4-channel surface coil. Localization and anatomical reference were achieved using T2 weighted Turbo Spin Echo Sequences in all three geometrical orientations (axial, coronal, longitudinal). The same acquisition field-of-view (FOV) and geometric parameters, including the number of slices to cover the whole brain (10-11), the slice thickness (0.5-1 mm), and the in-plane spatial resolution (∼273 µm) were used for all 2D scans (including DCE-MRI) to facilitate post-acquisition processing. The following sequences were used to obtain our MRI data: 1) gradient-echo sequences (GRE) with TR =100 msec and TE = 2 msec and multiple flip-angles(FA=5,10,15,30,45,70,80,90) to acquire T1 maps of the brain before injection of contrast; 2) a GRE sequence with TR = ∼50 msec and TE = ∼2 msec repeated 100 times with a temporal resolution of ∼3.6 sec to obtain the DCE-MRI data. The injection of MR contrast was done as a bolus through a tail vein catheter around the 10^th^ volume in order to follow the agent’s vascular kinetics; 3) a 3D GRE sequence was then used to obtain images with 110 µm isotropic resolution. Following the transfer of DICOM images, post-processing, extraction of T1 maps, and DCE-MRI processing was done using a built-in tool included in JIM 7.0 imaging software (Xinapse Systems). DCE-MRI processing involved using the T1 maps to obtain concentration from the MR image sequence and then using an automated arterial input function algorithm to calibrate the signal from different regions and tissues as described by Xinape Systems.^28^ A model-independent voxel-by-voxel parametric map was then generated depicting the integrated area under the curve (IAUC) at different times after injection. The IAUC at 30 sec and 120 sec were selected to provide a semi-quantitative non-specific measure of perfusion and permeability. Delineation of tumors in 2D and 3D MR images and parametric maps extracted from the DCE-MRI sequence was done using semi-automated threshold-based segmentation approaches included in ITK_SNAPsoftware. Tumor volumes were then obtained and expressed in mm_3_.

### Statistical Analysis

All statistical analyses were performed using Graphpad Prism 8 (GraphPad, San Diego, CA). The sample size for each group was ≥ 3, and data were reported as mean ± SD. Multiple comparisons were done using one-way ANOVA. Comparisons between two groups were done using the Student t-test. All *p* values were considered statistically significant if <0.05, and labeled as * p<0.05, ** p<0.01, and *** p<0.001. A Kaplan-Meier survival curve was generated to record animal survival, and a log-rank test was applied to compare survival between control and irradiated mice.

## Results

### 5F7 Binding to HER2 expressing BT474Br cells in vitro and in vivo

Staining of BT474Br cells *in vitro* with 5F7 and detection using anti-alpaca-biotinylated antibody and streptavidin-FITC revealed a positive signal compared to negative controls (Fig. 1A). Flow cytometry analysis of 5F7 binding to live BT474Br cells incubated with and counter-stained with anti-alpaca-biotin and streptavidin-FITC indicated that about 97% of cells were positive for HER2 expression (Fig. 1B, C). The staining of BT474Br tumor tissue sections from mouse brain for HER2 with 5F7 further validated its binding to BT474Br cells *ex vivo* (Fig. 1D, F, G). No binding of 5F7 was detected in normal brain tissue (Fig. 1E, G), further validating the specificity of 5F7 binding to HER2-expressing BCBM.

**Fig. 1.**
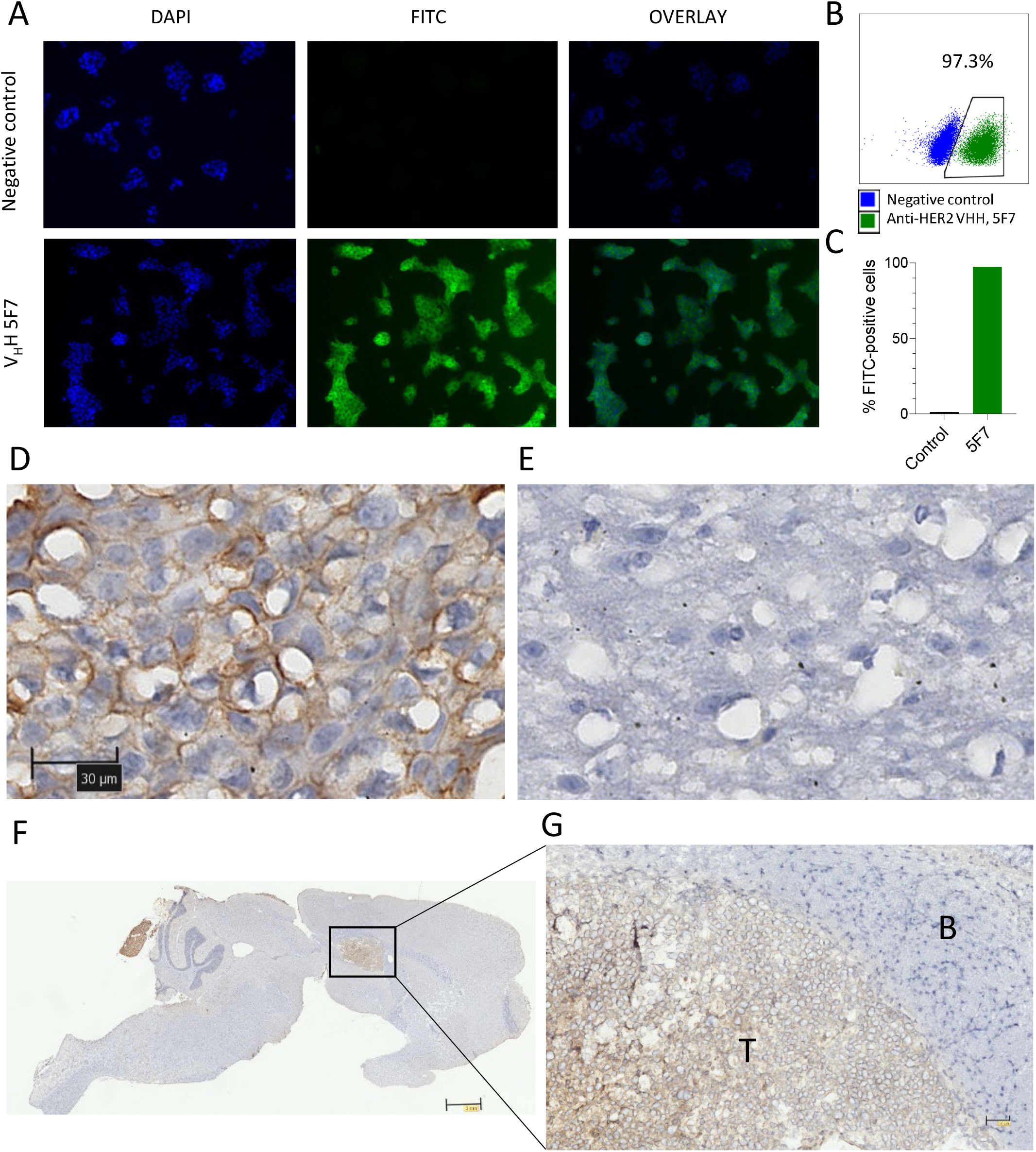
5F7 binding to HER2-positive breast cancer brain metastasis cells. (A) Staining of BT474Br cells with 5F7 and detection using anti-alpaca-biotinylated antibody and streptavidin-FITC. (B) Flow cytometry analysis of 5F7 binding to live BT474BrM3 cells incubated with and counter-stained with anti-alpaca-biotin and streptavidin-FITC. (C) Quantitative analysis demonstrated that 94% of cells express HER2. Staining with secondary antibody served as a negative control. (D) H&E staining of BT474BrM3 tumor tissue section from mouse brain for HER2 with 5F7 (10 μg/ml) and biotinylated anti-alpaca antibody. Detection of secondary antibodies with streptavidin-peroxidase. The scale bar is 30-μm (E) Negative staining of normal brain tissue from the frontal lobe. (F) 5F7 bound to tumor cells after systemic delivery was detected using an anti-alpaca-biotinylated antibody and streptavidin-peroxidase. Specific stain seen in tumor but not in normal brain. Scale bar is 2 mm. (G) A higher power magnification of region from image F. T-tumor, B-normal brain. The scale bar is 100 μm.

### _18_F-5F7 Uptake In Intracranial BT474Br Xenografts

Radiochemical yield for the conjugation of the prosthetic agent [^18^F]RL-III to 5F7 deriving _18_F-5F7 was 46.3 ± 7.7% and the molar activity of ^18^F-5F7 was 6.9 ± 4.9 GBq/µmol. The effects of irradiation on the uptake of ^18^F-5F7 was evaluated by longitudinal PET imaging in two cohorts of mice – the first imaged on Days 3 and 8 post-irradiation, and the second on Days 12 and 18 post-irradiation. No significant difference in tumor uptake between control and irradiated mice was observed until Day 12 (irradiated, 2.18 ± 1.18 %ID/g; control. 0.00 ± 0.00 %ID/g, **p<0.01) (Fig. 2A, B, D, E). On Day 18, irradiated mice also displayed a greater tumor uptake (2.54 ± 1.53 %ID/g, *p<0.05) than controls (0.26 ± 0.58 %ID/g) (Supplemental Table 1). Plotting ^18^F-5F7 uptake versus tumor volume revealed similar deviation from zero in slopes between control (p=0.013) and irradiated groups on Day 8 (p=0.051), with a significant slope difference being observed in the irradiated group (deviation from zero, p=0.012) on day 18 (Fig. 2C, F).

**Fig. 2.**
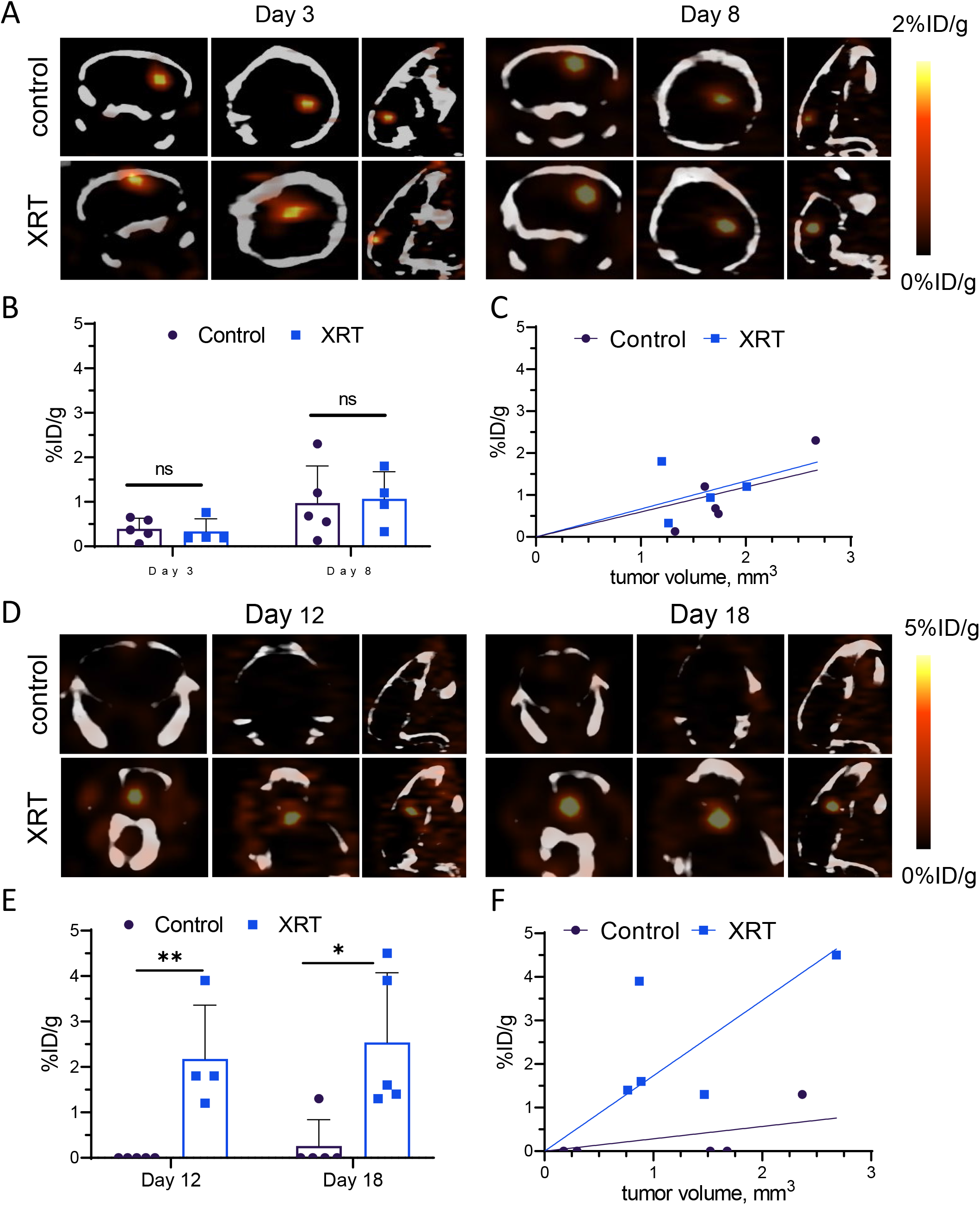
MicroPET/CT imaging of ^18^F-5F7 distribution in mice with intracranial BT474Br xenografts. Cohort 1: Days 3 and 8 post irradiation. (A) MicroPET/CT scans of representative control (n=5) and irradiated mice (n=4). (B) _18_F-5F7 uptake (% ID/g). (C) _18_F-5F7 uptake vs. tumor size (determined by morphometry on tissue sections). Cohort 2: Days 12 and 18 post irradiation. D. MicroPET/CT scans of representative control (n=5) and irradiated mice (n=4). (E). _18_F-5F7 uptake (% ID/g). F. [_18_F]5F7 uptake vs. tumor volume. Data presented as mean ± SD, * *P*<0.05; ** *P*<0.01. Unaired t-test.

### HER2 Expression in BT474Br Xenografts: Control vs. Irradiated Mice

We next investigated whether differences in _18_F-5F7 tumor uptake between control and irradiated mice was due to altered HER2 expression. After the last PET imaging session on Day 18, tissue sections from paraffin-embedded brains (n=3 from both irradiated and control groups) were obtained and evaluated for HER2 expression. No qualitative differences in HER2 expression were observed between control and irradiated BT474Br xenografts (Fig. 3A). Western blot analysis of cells from the brain tumors obtained two days after irradiation (n=4) and corresponding controls was performed to confirm these results; no difference in total HER2 expression was detected in BT474Br tissue from control and irradiated groups (p>0.05) (Fig. 3B). Similarly, flow cytometry showed no difference in HER2 expression on BT474Br cells harvested from the brains of control and irradiated mice (Fig. 3C). Moreover, there was no difference in the mean fluorescence intensity of bound antibodies between BT474Br cells harvested from control and irradiated animals. These data confirm that WBRT does not change HER2 expression in intracranial BT474Br xenografts.

**Fig. 3.**
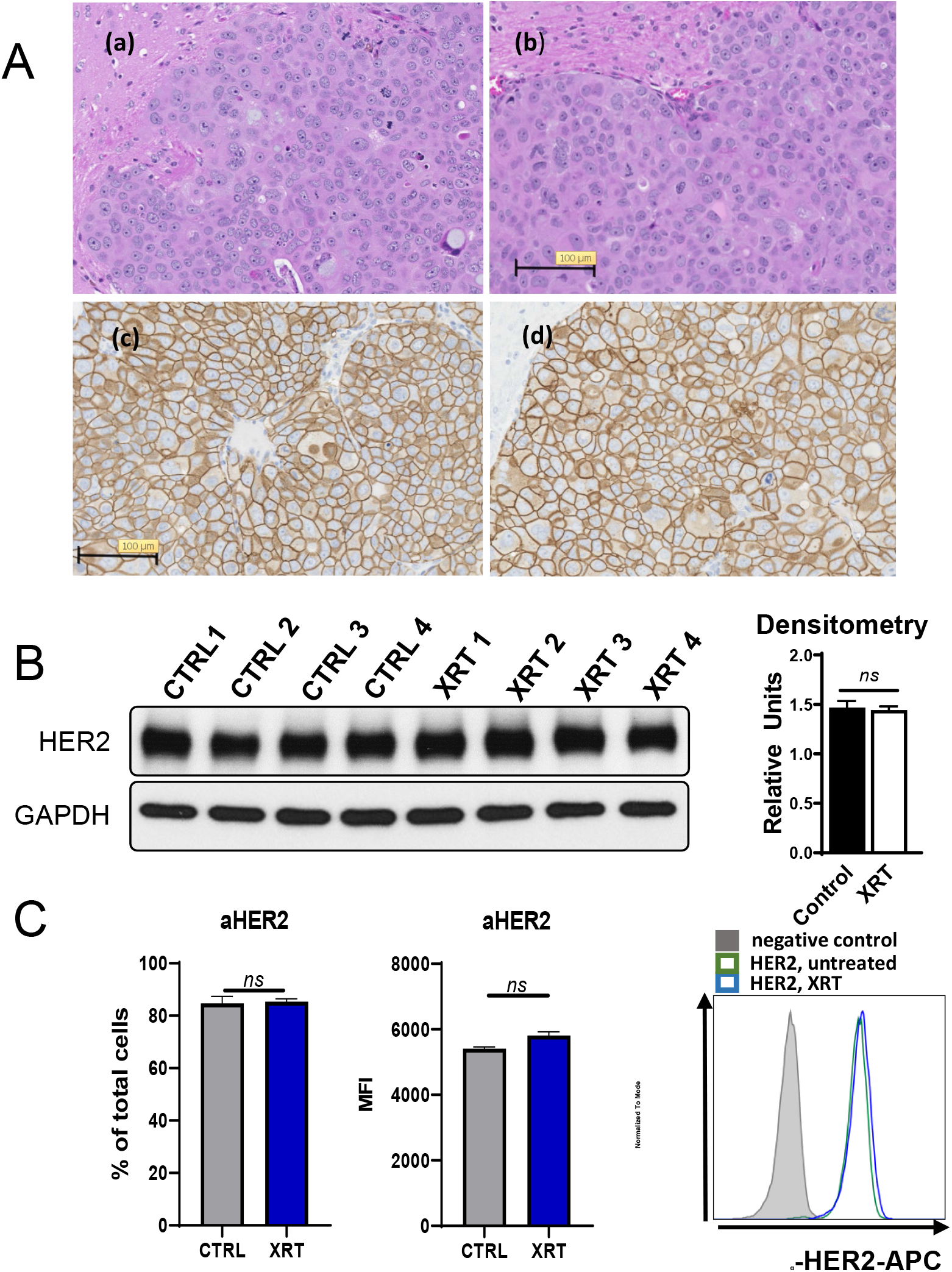
Analysis of HER2 expression in BCBM BT474Br model after WBRT by three complementary approaches -western blot, flow cytometry, and immunohistochemistry. (A) H&E stain of a representative brain section from control (a) and irradiated mice (b) and corresponding immunostaining for HER2 expression from control (c) and irradiated (d) mice. The scale bar is 100 µm. (B) HER2 expression in cells harvested from BT474Br tumors after 5 × 2 Gy daily WBRT and controls by western blot analysis. Densitometry of HER2 and GAPDH performed and calculated as a ratio show no significant difference between HER2 expression in control and irradiated animals. (n=4/group; mean ± SD, *P* > 0.05, Student’s t-test). (C) HER2-expression analysis on BT474Br cells harvested from control and irradiated tumors by flow cytometry. Results expressed as % positive cells and median fluorescent intensity (MFI). A representative flow histogram is shown on the right (negative control stain is grey, stains for HER2 expression of control and irradiated cells are blue and green, respectively). Data presented as mean ± SD, *P*>0.05; ns-not significant. Student’s t-test.

### MRI Evaluation of BT474Br Tumor Volume and Vascular Status

MRI-based BT474Br tumor volumetric analysis was performed to evaluate temporal changes in tumor volume in response to WBRT. Fig. 4A (a) shows a representative slice of a mouse brain from the 3D MRI data set with the tumor region depicted in red color. The corresponding rendered 3D images (b,c) depict two views of the head/brain outline and 3D tumor morphology, illustrating the ability to visualize the location, size, and morphology of the tumor lesion. The average tumor volume at 3, 7, and 17 days after irradiation did not change significantly in the treated group; however, there was a significant increase in size in the control group on day 17 (Fig. 4B). Although MRI indicated a difference in tumor volume between control and irradiated mice on day 17, there was no difference in median survival between these two groups (Fig. 4C). Representative control and irradiated mouse MR images with superimposed parametric IAUC maps extracted from the DCE-MRI data are shown in Fig. 5A. The longitudinal parametric changes in tumor perfusion/permeability compared to normal brain consistently exhibited a markedly more heterogeneous and often larger increase in both IAUC30 and IAUC120 in the irradiated group compared to the controls, especially at 7 days post-irradiation (Fig. 5B, C). Tumor IAUC values were consistently higher than for normal brains in both treated and control animals. The longitudinal changes observed in the irradiated mice suggest more heterogeneous intra-tumoral micro-environment alterations than in the control mice. The longitudinal changes observed in tumor IAUC30 were statistically different between the two cohorts (compared to the pre-radiation baseline value), but the IAUC120 were not, although an increasing trend was seen. Although the effect of radiation on tumor vasculature parameters was consistently observed, a comparison of average whole-tumor values fails to capture the complex changes induced. This is illustrated in the 2D, and the corresponding 3D Intratumor vascular variability maps (Fig. 6), demonstrating more heterogeneous behavior in irradiated mice, particularly in the 3D rendered images.

**Fig. 4.**
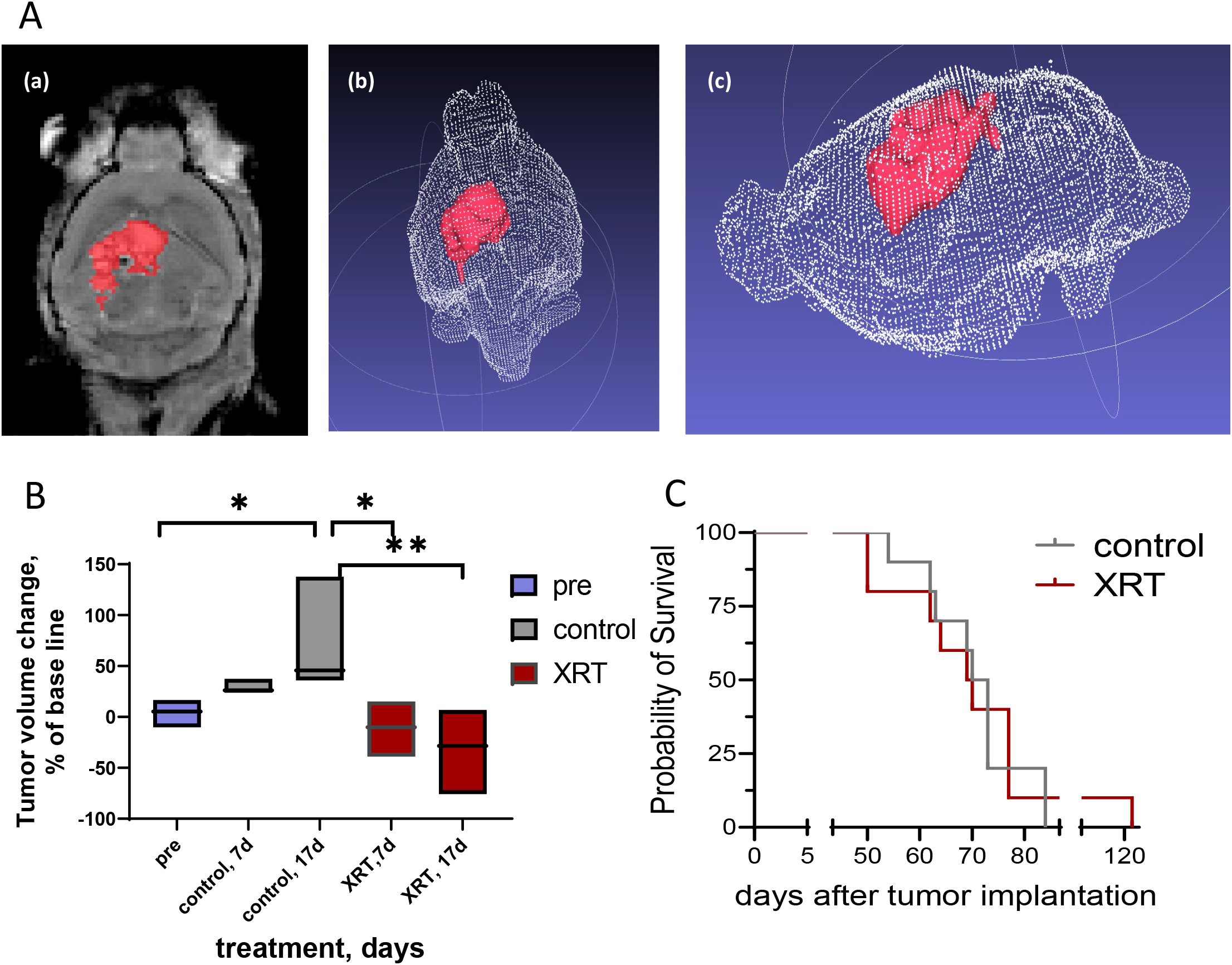
Effect of WBRT on tumor volume and survival. (A) (a) T2 MRI visualizes BT474Br intracranial tumor (red) in 2D (a) and two 3D (b,c) views. (B) Tumor volume changes post WBRT compared to untreated controls. (C). Survival analysis shows no significant difference between irradiated and control groups. Data presented as mean ± SD, * *P*<0.05; ** *P*<0.01. One-way ANOVA, Tukey’s multiple comparisons test.

**Fig. 5.**
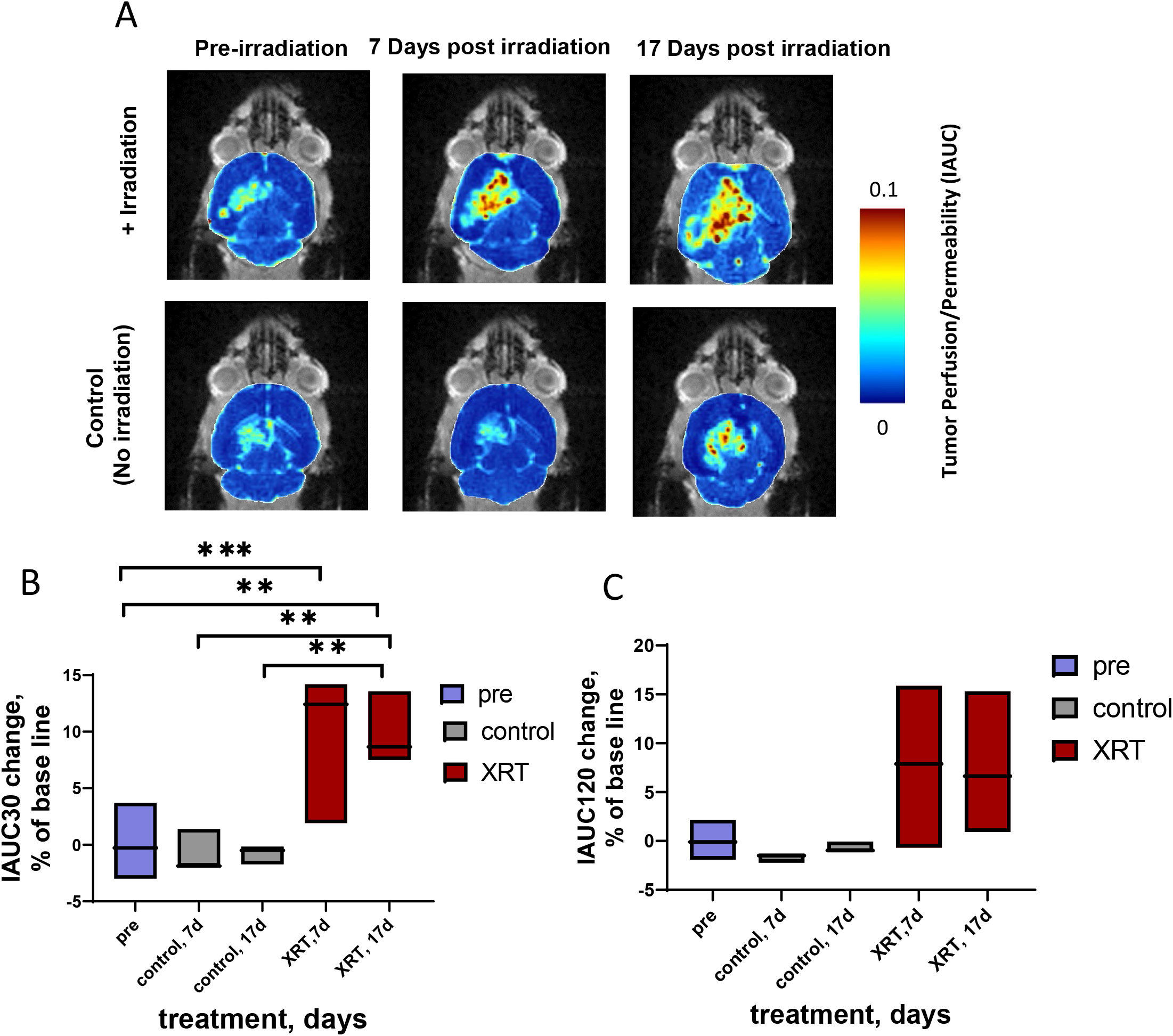
Tumor perfusion and permeability visualization, and IAUC30 and IAUC120 parameters from DCE-MRI data. (A) IAUC parametric color maps of the brain and tumor of representative mice from irradiated and control groups superimposed on anatomical MRI for reference. Vascular changes in tumors and normal brain due to WBRT are readily visualized. The average percent short (B; IAUC 30) and long (C; IAUC 120)-time uptake and washout kinetics of contrast were determined in mice before (baseline) and at 7and 17 days post-irradiation. * *P*<0.05; ** *P*<0.01; *** *P*<0.001. One-way ANOVA, Tukey’s multiple comparisons test.

**Fig. 6.**
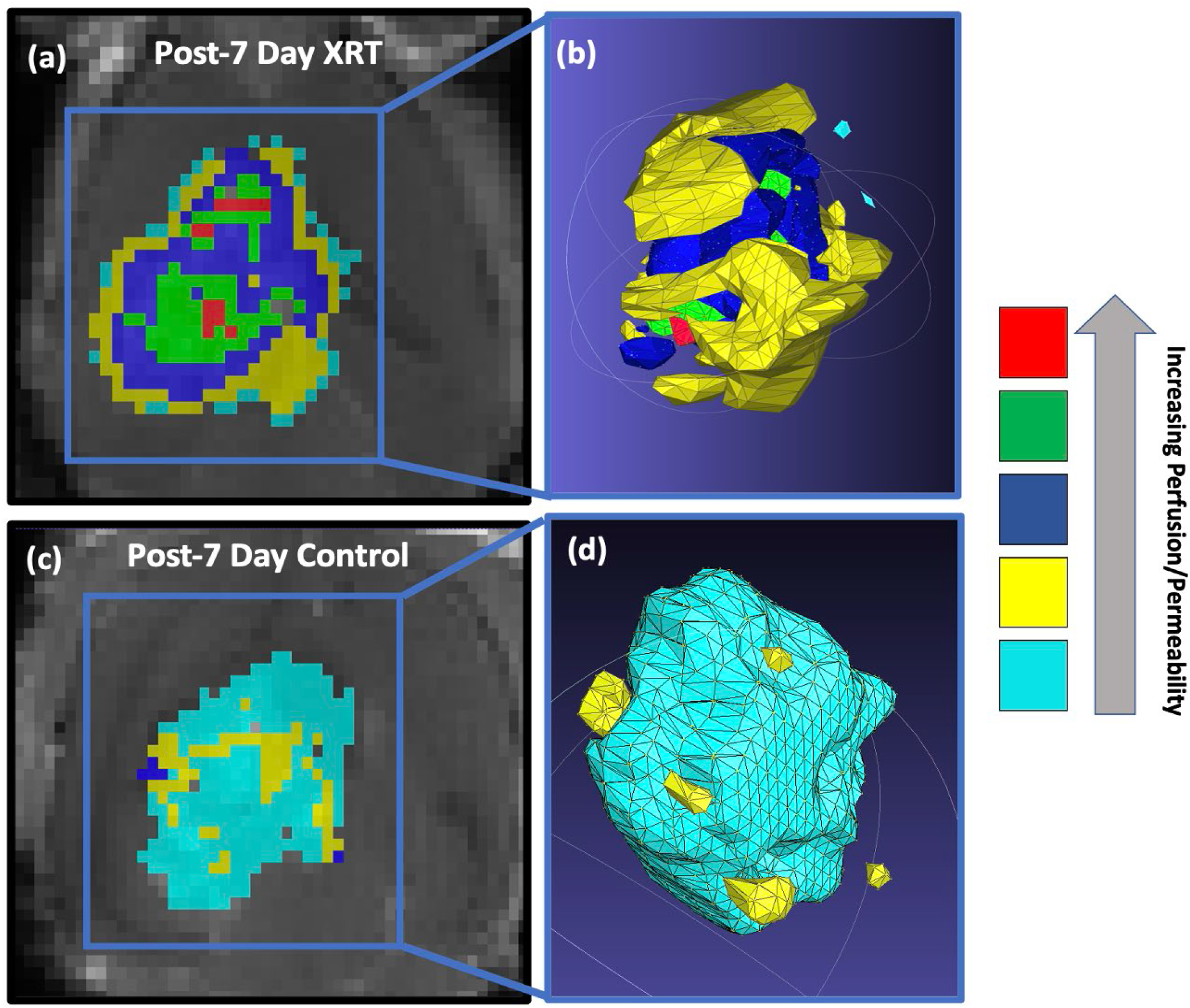
Visualization of intra-tumor perfusion and permeability heterogeneity on Day 7. IAUC120 derived color maps quantitatively segmented and superimposed on the corresponding MRI anatomical images for representative irradiated (a) and control (c) mouse illustrating intra-tumoral patterns of perfusion/permeability. The corresponding 3D renderings (b, irradiated; d, control) enable 3D visualization of radiation-induced heterogeneities in the tumor microenvironment.

## DISCUSSION

Treatment of brain metastases is challenging due to the need to spare healthy normal brain tissue while achieving adequate drug transport across the BTB to have sufficient therapeutic effect. HER2 targeting via V_H_H molecules offers a promising approach for targeting radiation to HER2 positive cancers, as demonstrated by studies with 5F7 labeled with both the therapeutic radionuclides ^131^I^22^ and ^211^At^29^ and ^18^F for PET imaging.^20^ Likewise, radiolabeled V_H_H tumor targeting of therapeutic and PET radionuclides also has been demonstrated with 2Rs15d, an anti-HER2 V_H_H, which binds to a different epitope on the HER2 extracellular domain than 5F7^20,30^ and has recently entered clinical investigation.^31^ Studies with radiolabeled 2Rs15d confirmed the ability to target intracranial tumors in mice;^32^ however, the therapeutic effect was modest even with 3 doses of ^225^Ac-2Rs15d, which emits 4 α-particles per decay.

In order to evaluate whether low-dose WBRT could augment V_H_H uptake in intracranial xenografts, we utilized the HER2-positive brain metastatic breast cancer line, BT474Br.^33,34^ We first validated that 5F7 efficiently binds to HER2 on BT474Br cells *in vitro* by immunocytochemistry and flow cytometry. Positive and specific detection of HER2-expressing BT474Br cells in xenografts but not normal brain also was evident on tissue sections. Furthermore, 5F7 accumulated in tumor tissue but not in the normal brain after i.v. delivery, confirming that this V_H_H is suitable for imaging HER2-positive BCBM.

We selected WBRT as a strategy for enhancing the potential utility of radiolabeled V_H_H for the detection and treatment of BCBM. Patients undergoing WBRT for treatment of brain metastases typically receive a fractionated 58–60 Gy regimen, with approximately 10% surviving beyond 12 months.^35^ In our experiments, mice were subjected to a fractionated WBRT of 10 Gy (5 × 2 Gy daily) regimen to evaluate a dose likely avoiding neurotoxicity in patients, limiting the value of WBRT for patients with BCBM at higher doses.^36,37^ In addition, we employed longitudinal PET imaging with ^18^F-5F7 to quantify tumor delivery noninvasively in the same animals at multiple time points after irradiation. As expected, at this radiation dose, WBRT did not improve the survival of the mice. However, we observed a significant increase in ^18^F-5F7 V_H_H uptake beginning 12 days after WBRT completion compared to non-irradiated mice. Furthermore, the enhanced uptake of the 5F7 V_H_H was more pronounced on day 18 after WBRT motivating future studies to define the post-WBRT time when uptake enhancement would occur.

Radiotherapy can change gene expression in several cell types within a tumor and a normal brain.^38,39^ To ensure that the increase in ^18^F-5F7 uptake was not due to WBRT-induced alterations in HER2 expression, we analyzed the HER2 levels in tumor tissue ex vivo obtained after the PET imaging experiment. H&E staining of tissue sections confirmed tumor presence in both irradiated and control mice. The measurement of HER2 expression levels revealed no difference between groups, confirming that the higher uptake of ^18^F-5F7 in the tumors of irradiated mice was not due to changes in HER2 expression. Consistent with the well-known effects of radiation on tumor vasculature,^40-42^ it is likely that the higher uptake of ^18^F-5F7 in irradiated tumors reflects their higher permeability.

To gain further insights into the effects of WBRT on ^18^F-5F7 uptake in intracranial BT474Br xenograft, we utilized DCE-MRI with Gd^3+^ contrast to assess BTB permeability in irradiated and control mice. MRI provided a volumetric comparison of tumors, which demonstrated a significant increase in tumors’ growth in control but not WBRT groups at day 17 but not at earlier time points. In contrast, we observed greater tumor perfusion and permeability of contrast agent in irradiated tumors with DCE-MRI than in non-irradiated controls. The difference in permeability between control and irradiated animals was apparent about a week after treatment, with more distinction on Day 17 after WBRT. The changes in the permeability of BT474Br BCBM after irradiation correlated with a greater uptake of ^18^F-5F7, particularly at the latest time points analyzed in our study. Our data are consistent with a previous report where patients with brain metastases had an increase in gefitinib uptake with increasing doses of WBRT up to 30 Gy.^43^ In addition, several studies in preclinical mouse models have found that BCBM lesion volume does not correlate with passive permeability.^13,44,45^

The number of lesions in BCBM patients can vary significantly from a single lesion to greater than 10, which is a factor in determining a treatment regimen. WBRT, the most prevalent treatment, is performed at different intervals to mitigate the associated neurocognitive decline.^46^ In a 30-patient study, the permeability of metastatic brain lesions was found to be variable and could be increased to varying degrees about 2-4 weeks after WBRT or stereotactic radiosurgery,^47^ Although not investigated in depth in this study, our DCE-MRI imaging revealed heterogeneity in the perfusion/vascular permeability alterations induced by the irradiation of BT474Br tumors.

Nevertheless, by increasing the permeability of BCBM and addressing the heterogeneous response of tumor vasculature to irradiation, we may unlock the potential for administering more targeted therapies to these patients. This could avoid the need for WBRT at doses causing unwanted side effects and compromising quality of life. Considering the type of vehicle for tumor targeting, the permeability requirements for V_H_H should be favorable due to their considerably smaller size compared to intact antibodies.

Although WBRT is known to enhance the permeability of the brain-tumor-barrier in both animal models and patients, its effects have not yet been validated for use with small protein molecules like V_H_H. Studies in animal models have demonstrated that the BTB in brain metastases is more restrictive than in glioblastoma,^48^ making V_H_H a relevant carrier system for theranostic approaches for brain metastases. Future work will explore these requirements as we look to define the optimal XRT regimen for increasing permeability without inflicting irreversible damage to the normal brain.

Overcoming the BTB would allow new diagnostic tools to detect metastases that would otherwise remain undiscovered with existing strategies. In addition, a greater diagnostic toolbox would better inform the selection of treatment regimens. In conclusion, our V_H_H PET tracer, ^18^F-5F7, exhibited enhanced tumor accumulation in a HER2-expressing BCBM mouse model when preceded by low-dose WBRT. The increased uptake of 5F7 V_H_H after WBRT in tumors correlated with increased perfusion and permeability as measured by the DCE-MRI and was not linked to changes in HER2 expression. These studies suggest the potential utility of the V_H_H PET tracer, ^18^F-5F7, as an imaging tool for HER2-positive BCBM. The novelty lies in the ability of 5F7 V_H_H to target brain lesions and not normal brain after WBRT and the added opportunity to use DCE-MRI to visualize and quantify changes in vascular parameters in this context. The knowledge we gained in this study could be optimized for therapeutic intervention to increase the uptake of V_H_H carrier molecules labeled for imaging and potentially therapeutic radionuclides.^49^ These results bring us closer to understanding how BTB permeability can be increased to deliver small proteins like V_H_H to BCBM and warrant further investigation.

## ACKNOWLEDGMENTS

The authors are grateful to the Mouse Histology and Phenotyping Core Facility, Northwestern University, for assistance with processing and staining mouse tissues.

## REFERENCES

1. Howlader N, Noone AN, Krapcho M, et al. (eds). SEER Cancer Statistics Review, 1975–2018. National Cancer Institute. Bethesda, MD..

2. Savci-Heijink CD, Halfwerk H, Hooijer GKJ, Horlings HM, Wesseling J, van de Vijver MJ. Retrospective analysis of metastatic behaviour of breast cancer subtypes. Breast Cancer Res Treatment. 2015;150:547–557.

3. Giordano SH, Buzdar AU, Smith TL, Kau SW, Yang Y, Hortobagyi GN. Is Breast Cancer Survival Improving? Trends in Survival for Patients with Recurrent Breast Cancer Diagnosed from 1974 through 2000. Cancer. 2004;100:44–52.

4. Haffty BG, Yang Q, Reiss M, et al. Locoregional relapse and distant metastasis in conservatively managed triple negative early-stage breast cancer. J Clin Oncol. 2006;24:5652–5657.

5. Bauer KR, Brown M, Cress RD, Parise CA, Caggiano V. Descriptive analysis of estrogen receptor (ER)-negative, progesterone receptor (PR)-negative, and HER2-negative invasive breast cancer, the so-called triple-negative phenotype: A population-based study from the California Cancer Registry. Cancer. 2007;109:1721–1728.

6. Dawood S, Broglio K, Gonzalez-Angulo AM, Buzdar AU, Hortobagyi GN, Giordano SH. Trends in survival over the past two decades among White and Black patients with newly diagnosed stage IV breast cancer. J Clin Oncol. 2008;26:4891–4898.

7. Gil-Gil MJ, Martinez-Garcia M, Sierra A, et al. Breast cancer brain metastases: a review of the literature and a current multidisciplinary management guideline. Clin Trans Oncol. 2014;16:436–446.

8. Hayes DF, Thor AD, Dressler LG, et al. HER2 and Response to Paclitaxel in Node-Positive Breast Cancer. NEJM. 2007;357:1496–1506.

9. Moasser MM. The oncogene HER2: Its signaling and transforming functions and its role in human cancer pathogenesis. Oncogene. 2007;26:6469–6487.

10. Hurvitz SA, O’Shaughnessy J, Mason G, et al. Central Nervous System Metastasis in Patients with HER2-Positive Metastatic Breast Cancer: Patient Characteristics, Treatment, and Survival from SystHERs. Clin Cancer Res. 2019;25(8):2433–2441.

11. Leyland-Jones B. Human Epidermal Growth Factor Receptor 2–Positive Breast Cancer and Central Nervous System Metastases. J Clin Oncol. 2009;27:5278–5286.

12. Niwinska A, Murawska M, Pogoda K. Breast cancer brain metastases: differences in survival depending on biological subtype, RPA RTOG prognostic class and systemic treatment after whole-brain radiotherapy (WBRT). Annals Oncol. 2010;21:942–948.

13. Lockman PR, Mittapalli RK, Taskar KS, et al. Heterogeneous Blood–Tumor Barrier Permeability Determines Drug Efficacy in Experimental Brain Metastases of Breast Cancer. Clin Cancer Res. 2010;16:5664–5678.

14. DeAngelis LM, Delattre J-Y, Posner JB. Radiation-induced dementia in patients cured of brain metastases. Neurology. 1989;39:789–789.

15. Patchell RA. The management of brain metastases. Cancer Treat Rev.2003;29:533–540.

16. Jakubovic R, Sahgal A, Soliman H, et al. Magnetic Resonance Imaging-based Tumour Perfusion Parameters are Biomarkers Predicting Response after Radiation to Brain Metastases. Clin Oncol. 2014;26:704–712.

17. Dijkers EC, Oude Munnink TH, Kosterink JG, et al. Biodistribution of 89Zr-trastuzumab and PET imaging of HER2-positive lesions in patients with metastatic breast cancer. Clin Pharmacol Ther. 2010;87(5):586–592.

18. Debie P, Lafont C, Defrise M, et al. Size and affinity kinetics of nanobodies influence targeting and penetration of solid tumours. J Control Release. 2020; 317:34–42.

19. Altunay B, Morgenroth A, Beheshti M, et al. HER2-directed antibodies, affibodies and nanobodies as drug-delivery vehicles in breast cancer with a specific focus on radioimmunotherapy and radioimmunoimaging. Eur J Nucl Med Mol Imaging. 2021;48(5):1371–1389.

20. Zhou Z, Vaidyanathan G, McDougald D, et al. Fluorine-18 Labeling of the HER2-Targeting Single-Domain Antibody 2Rs15d Using a Residualizing Label and Preclinical Evaluation. Mol Imag Biol. 2017;19:867–877.

21. Keyaerts M, Xavier C, Heemskerk J, et al. Phase I Study of 68Ga-HER2-Nanobody for PET/CT Assessment of HER2 Expression in Breast Carcinoma. J Nucl Med. 2016;57:27–33.

22. Pruszynski M, Koumarianou E, Vaidyanathan G, et al. Targeting breast carcinoma with radioiodinated anti-HER2 Nanobody. Nucl Med Biol. 2013;40:52–59.

23. Shaw MG, Ball DL. Treatment of Brain Metastases in Lung Cancer: Strategies to Avoid/Reduce Late Complications of Whole Brain Radiation Therapy. Current Treat Options Oncol. 2013;14:553–567.

24. Zhang S, Huang W-cC, Zhang L, et al. Src family kinases as novel therapeutic targets to treat breast cancer brain metastases. Cancer Res. 2013;73:5764–5774.

25. Balyasnikova IV, Prasol MS, Ferguson SD, et al. Intranasal Delivery of Mesenchymal Stem Cells Significantly Extends Survival of Irradiated Mice with Experimental Brain Tumors. Molecular Therapy. 2014;22:140–148.

26. Dey M, Yu D, Kanojia D, et al. Intranasal Oncolytic Virotherapy with CXCR4-Enhanced Stem Cells Extends Survival in Mouse Model of Glioma. Stem Cell Reports. 2016;7:471–482.

27. Zhou Z, McDougald D, Meshaw R, Balyasnikova I, Zalutsky MR, Vaidyanathan G. Labeling single domain antibody fragments with (18)F using a novel residualizing prosthetic agent - N-succinimidyl 3-(1-(2-(2-(2-(2-[(18)F]fluoroethoxy)ethoxy)ethoxy)ethyl)-1H-1,2,3-triazol-4-yl)-5-(guanidinomethyl)benzoate. Nucl Med Biol. 2021;100-101:24–35.

28. Shi L, Wang D, Liu W, et al. Automatic detection of arterial input function in dynamic contrast enhanced MRI based on affinity propagation clustering. J Magn Reson Imaging. 2014;39(5):1327–1337.

29. Choi J, Vaidyanathan G, Koumarianou E, Kang CM, Zalutsky MR. Astatine-211 labeled anti-HER2 5F7 single domain antibody fragment conjugates: radiolabeling and preliminary evaluation. Nucl Med Biol. 2018;56:10–20.

30. D’Huyvetter M, De Vos J, Xavier C, et al. 131 I-labeled Anti-HER2 Camelid sdAb as a Theranostic Tool in Cancer Treatment. Clinical Cancer Res. 2017;23:6616–6628.

31. D’Huyvetter M, Vos J, Caveliers V, et al. Phase I Trial of (131)I-GMIB-Anti-HER2-VHH1, a New Promising Candidate for HER2-Targeted Radionuclide Therapy in Breast Cancer Patients. J Nucl Med. 2021;62(8):1097–1105.

32. Puttemans J, Dekempeneer Y, Eersels JL, et al. Preclinical Targeted alpha- and beta(-)-Radionuclide Therapy in HER2-Positive Brain Metastasis Using Camelid Single-Domain Antibodies. Cancers (Basel). 2020;12(4).

33. Nanni P, Nicoletti G, Palladini A, et al. Multiorgan metastasis of human HER-2+ breast cancer in Rag2-/-;Il2rg-/-mice and treatment with PI3K inhibitor. PLoS ONE. 2012.;7

34. Palmieri D, Bronder JL, Herring JM, et al. Her-2 overexpression increases the metastatic outgrowth of breast cancer cells in the brain. Cancer Res. 2007;67:4190–4198.

35. Central nervous system tumours, 264–281(Elsevier 2011).

36. Brown PD, Jaeckle K, Ballman KV, et al. Effect of Radiosurgery Alone vs Radiosurgery With Whole Brain Radiation Therapy on Cognitive Function in Patients With 1 to 3 Brain Metastases: A Randomized Clinical Trial. JAMA. 2016; 316(4):401–409.

37. Chang EL, Wefel JS, Hess KR, et al. Neurocognition in patients with brain metastases treated with radiosurgery or radiosurgery plus whole-brain irradiation: a randomised controlled trial. Lancet Oncol. 2009;10(11):1037–1044.

38. Schulz M, Michels B, Niesel K, et al. Cellular and Molecular Changes of Brain Metastases-Associated Myeloid Cells during Disease Progression and Therapeutic Response. iScience. 2020;23(6):101178.

39. Svensson JP, Stalpers LJ, Esveldt-van Lange RE, et al. Analysis of gene expression using gene sets discriminates cancer patients with and without late radiation toxicity. PLoS Med. 2006;3(10):e422.

40. Dilworth JT, Krueger SA, Dabjan M, et al. Pulsed low-dose irradiation of orthotopic glioblastoma multiforme (GBM) in a pre-clinical model: effects on vascularization and tumor control. Radiother Oncol. 2013;108(1):149–154.

41. Yuan H, Gaber MW, Boyd K, Wilson CM, Kiani MF, Merchant TE. Effects of fractionated radiation on the brain vasculature in a murine model: blood-brain barrier permeability, astrocyte proliferation, and ultrastructural changes. Int J Radiat Oncol Biol Phys. 2006;66(3):860–866.

42. Warrington JP, Ashpole N, Csiszar A, Lee YW, Ungvari Z, Sonntag WE. Whole brain radiation-induced vascular cognitive impairment: mechanisms and implications. J Vasc Res. 2013;50(6):445–457.

43. Zeng Y-D, Liao H, Qin T, et al. Blood-brain barrier permeability of gefitinib in patients with brain metastases from non-small-cell lung cancer before and during whole brain radiation therapy. Oncotarget. 2015;6:8366–8376.

44. Adkins CE, Mohammad AS, Terrell-Hall TB, et al. Characterization of passive permeability at the blood–tumor barrier in five preclinical models of brain metastases of breast cancer. Clin Exper Met. 2016;33:373–383.

45. Murrell DH, Hamilton AM, Mallett CL, Gorkum RV, Chambers AF, Foster PJ. Understanding heterogeneity and permeability of brain metastases in murine models of HER2-positive breast cancer through magnetic resonance imaging: Implications for detection and therapy. Translational Oncol. 2015;8:176–184.

46. Leone JP, Leone BA. Breast cancer brain metastases: the last frontier. Exper Hematol Oncol. BioMed Central; 2015;4:33.

47. Teng F, Tsien CI, Lawrence TS, Cao Y. Blood-tumor barrier opening changes in brain metastases from pre to one-month post radiation therapy. Radiother Oncol. 2017;125:89–93.

48. Mittapalli RK, Adkins CE, Bohn KA, Mohammad AS, Lockman JA, Lockman PR. Quantitative Fluorescence Microscopy Measures Vascular Pore Size in Primary and Metastatic Brain Tumors. Cancer Res. 2017;77(2):238–246.

49. Feng Y, Meshaw R, McDougald D, et al. Evaluation of an (131)I-labeled HER2-specific single domain antibody fragment for the radiopharmaceutical therapy of HER2-expressing cancers. Sci Rep. 2022;12(1):3020.

