## Supplemental Table 1 for "Low Level Whole-Brain Radiation Enhances Theranostic Potential Of Single Domain Antibody Fragments For HER2-Positive Brain Metastases"

^#^These authors contributed equally to this project.

^*^Present address: Evergreen Theragnostics, Springfield NJ 07081 USA

**Supplemental Table 1. ^18^F-5F7 Uptake in Control and Irradiated Mice.**

| Day | Control Mice | | Irradiated Mice | |
| --- | --- | --- | --- | --- |
|  | % ID/g* | N | % ID/g* | N |
| 3 | 0.39 ± 0.24 | 5 | 0.34 ± 0.28 | 4 |
| 8 | 0.97 ± 0.83 | 5 | 1.07 ± 0.61 | 4 |
| 12 | 0.00 ± 0.00 | 5 | 2.18 ± 1.18 | 4 |
| 18 | 0.26 ± 0.58 | 5 | 2.54 ± 1.53 | 5 |

*Mean ± SD
